# Peptide from NSP7 is able to form amyloid-like fibrils: artifact or challenge to drug design?

**DOI:** 10.1101/2022.06.28.497929

**Authors:** Yuri Garmay, Aleksandr Rubel, Vladimir Egorov

**Affiliations:** Petersburg Nuclear Physics Institute named by B. P. Konstantinov of National Research Center, Kurchatov Institute, 1 mkr. Orlova roshcha, Gatchina 188300, Russia; Saint Petersburg State University, 7/9 Universitetskaya Emb., St Petersburg 199034, Russia; Smorodintsev Research Institute of Influenza, Russian Ministry of Health, 15/17 Ulitsa Prof. Popova, St. Petersburg 197376, Russia; Institute of Experimental Medicine, 12 Ulitsa Akademika Pavlova, St. Petersburg 197376, Russia

## Abstract

Using a set of programs for analyzing the primary structure of amyloidogenic proteins, we analyzed the primary structure of the of the COVID-19 NSP7 protein. Peptides containing potential amyloidogenic determinants in the primary structure have been synthesized and shown to be capable of forming amyloid-like fibrils *in vitro*. In the future, this peptide can be used as a basis for creating antiviral agents that act directly on the spatial structure of the NSP7 subunit of the COVID-19 polymerase.

## Introduction

One of the currently frequently discussed methods of antivirals design is the specific induction of changes in the structure and functionality of proteins by a prion-like mechanism [1]. Indeed, viral proteins tend to form amyloid-like fibrils [2], and at the same time, they often have low homology with human proteins. For most proteins, the induction of a conformational transition by peptides is a process that depends on the coincidence of the primary structure of the protein and the peptide acting on it [3], [4]. Together with the ability to amyloid chain reaction, this suggests that such peptides will be effective at low concentrations and, at the same time, will not affect host proteins. A number of peptides are known to have antiviral activity and act according to this mechanism [5]–[8]. The general strategy for the search for such drugs is the search for an amyloidogenic protein determinant and the creation of a peptide that carries this determinant. Such a peptide can specifically affect the parental protein at low concentrations, causing its conformational transition and loss of activity [9]. The peptide can be delivered to cells using carriers or in the form of RNA encoding it [10], which has the elements necessary for translation on the eukaryotic ribosome. At the same time, it must be kept in mind that the effect of amyloid-like viral structures on the body has not been studied enough [11], [12] however, for example, in the case of the influenza virus, the formation of amyloid-like fibrils by the PB1-F2 protein during the life cycle of the virus does not have a significant effect on the host organism [13]–[16].

The NSP7 protein is a processivity subunit of COVID19 RNA-dependent RNA polymerase [17]. In this work, we analyzed the primary structure of NSP7 and proposed sequences of the peptide capable of forming amyloid-like fibrils and having the potential to influence the conformation of this subunit through a prion-like mechanism.

## Materials and methods

### Analysis of the proteins primary structure

The amino acid sequences were analyzed using the ARCHES software [18], FoldAmyloid [19] and the original program for searching for mirror symmetry in protein sequences [20]. Protein sequence of NSP7 (P0DTD1) from the SWISSPROT database were used.

### Peptides PINP73

Peptides PINP73 (MVSLLSVLLSM, 1191.66 Da) and PINP74 (KLWAQCVQ, 975.17) were synthesized at OOO NPF VERTA, Russia, purity more than 80%. Peptides were dissolved in 5 μL of DMSO an then phosphate buffered saline (PBS) buffer was added to the 1 mg/ml peptide concentration (0.5% DMSO), and then incubated for 1 hour with agitation at orbital shaker (Eppendorf, USA) at 55 °C, 600 rpm.

### Atomic Force Microscopy

PINP73 peptide sample were diluted 70 times with water, after which 10 μL of solution were applied onto a freshly cleaved mica substrate. After 1 minute of incubation, the sample was dried in compressed air. Images (topography of the sample surface) were obtained in the semi-contact mode on an atomic force microscope “NT-MDT” (NT-MDT, Russia), with NSG01 probe. Image processing was performed using “Gwyddion” software [21].

### Thioflavin T assay

A modified ThT assay method by LeVine III [22] was used. Aliquots (30 μL) of peptide solutions were added to 970 μL of 10 mM ThT solution in PBS buffer. Measurements were performed in a HITACHI F-4010 fluorescence spectrophotometer (Hitachi, Japan) with excitation wavelength 440 nm (bandpass 5 nm) and emission wavelength 478 nm (bandpass 5 nm).

### Congo Red assay

Congo red dye (Sigma Aldrich, USA) at a concentration of 50 μM in PBS was mixed with the 10 μM peptide solution. As a control, a similar sample was used in which a buffer was added instead of the peptide. Absorption spectra were recorded on a BMG Clariostar (BMG, Germany), and difference spectrum was analyzed.

## Results and Discussion

We analyzed the primary structure of the NSP7 protein using three programs – Arches, which allows predicting the formation of hairpins characteristic of amyloid-like proteins [18], FoldAmyloid [19], which analyzes the local amino acid composition, and an original program for searching for mirror symmetry motifs [23]. The choice of these programs was due to the fact that in the course of our previous studies, the programs separately made it possible to predict amyloidogenic peptides [24][25][26]. Figure 1 shows the results of the search for potential amyloidogenic regions in NSP7.

**Figure 1.**
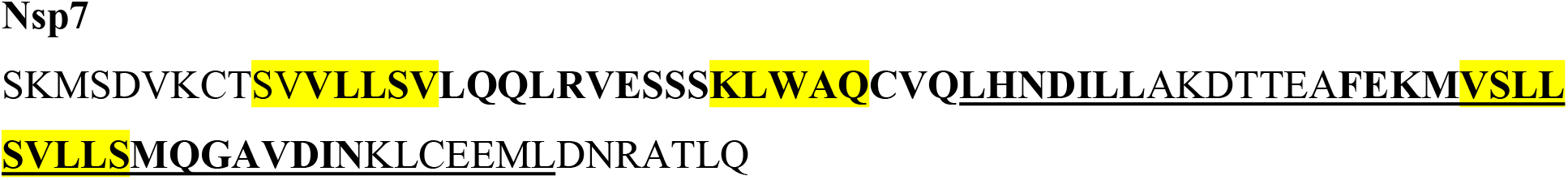
Amyloidogenic regions in NSP7 sequence. Symmetric part of protein is underlined, predicted with FoldAmyloid software amyloidogenic regions are colored yellow, predicted with Arches software are designated bold.

The peptide corresponding to the NSP7 sequence from 53 to 61 amino acid residues was determined as amyloidogenic by all three programs. A peptide corresponding to 52-62 residues (comprising 2 symmetrical methionines at the N- and C-terminus) was chemically synthesized, dissolved as described in Materials and Methods and analyzed using atomic force microscopy and two fluorescent dyes (Congo Red and Thioflavin T) characterizing amyloids. Atomic force microscopy topography image is shown on Figure 2. On the mica surface, filaments about 3 nm in height were observed, similar in morphology to amyloid-like fibrils.

**Figure 2.**
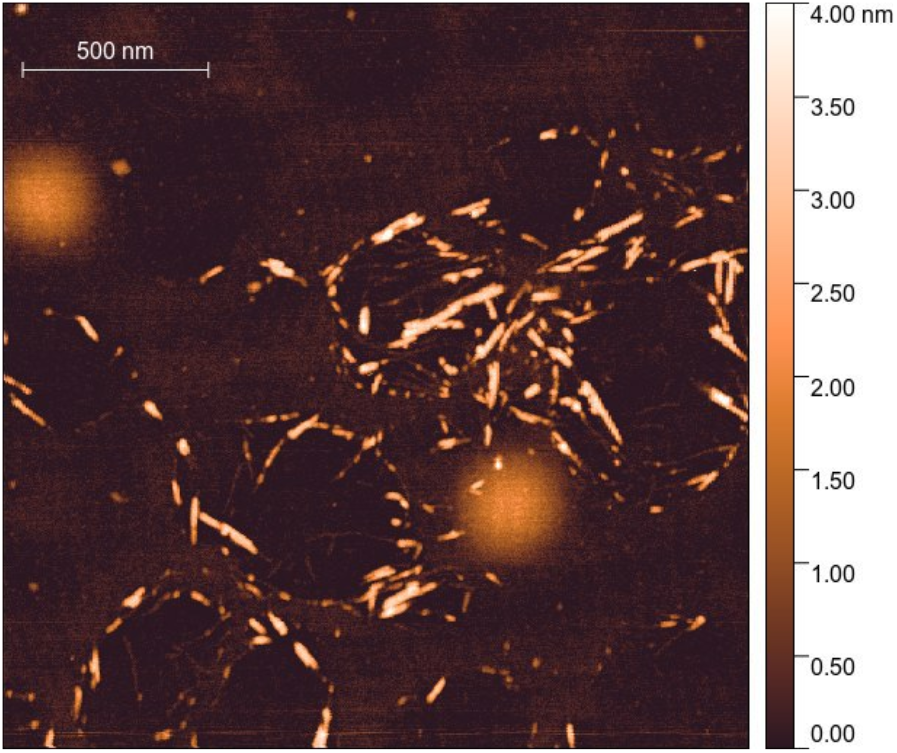
Atomic force microscopy topography image of PINP73 suspension on mica.

In order to determine the amyloidogenicity of the observed filaments, we performed fluorimetry in the presence of thioflavin T. As a control sample, we used a peptide PINP74, also identical to 27-37 amino acid residues NSP7 region, which has a similar molecular weight, but was determined as amyloidogenic only by FoldAmyloid.

An increase in thioflavin T fluorescence was observed in the presence of PINP73, but not PINP74 (Spectra are shown in Figure 3). Similarly, a comparison of the absorption spectrum of Congo red with PINP74 or PINP73 peptide showed the presence of a peak in the Congo Red with PINP73 spectrum in the region of 550 nm (Figure 4, peak is shown in difference spectrum).

**Figure 3.**
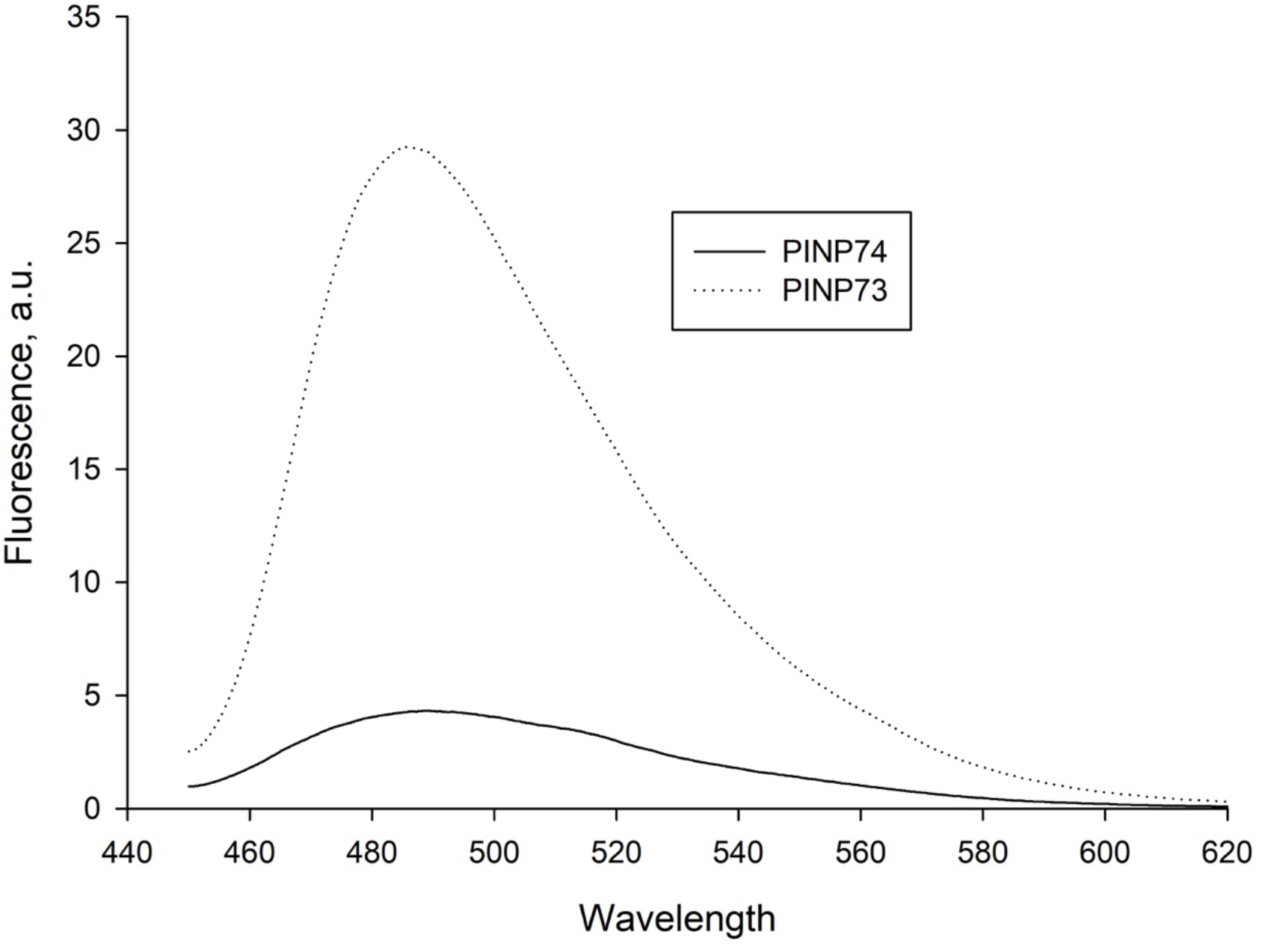
Fluorescence spectrum of PINP73 and PINP74 (control) solutions with Thioflavin T.

**Figure 4.**
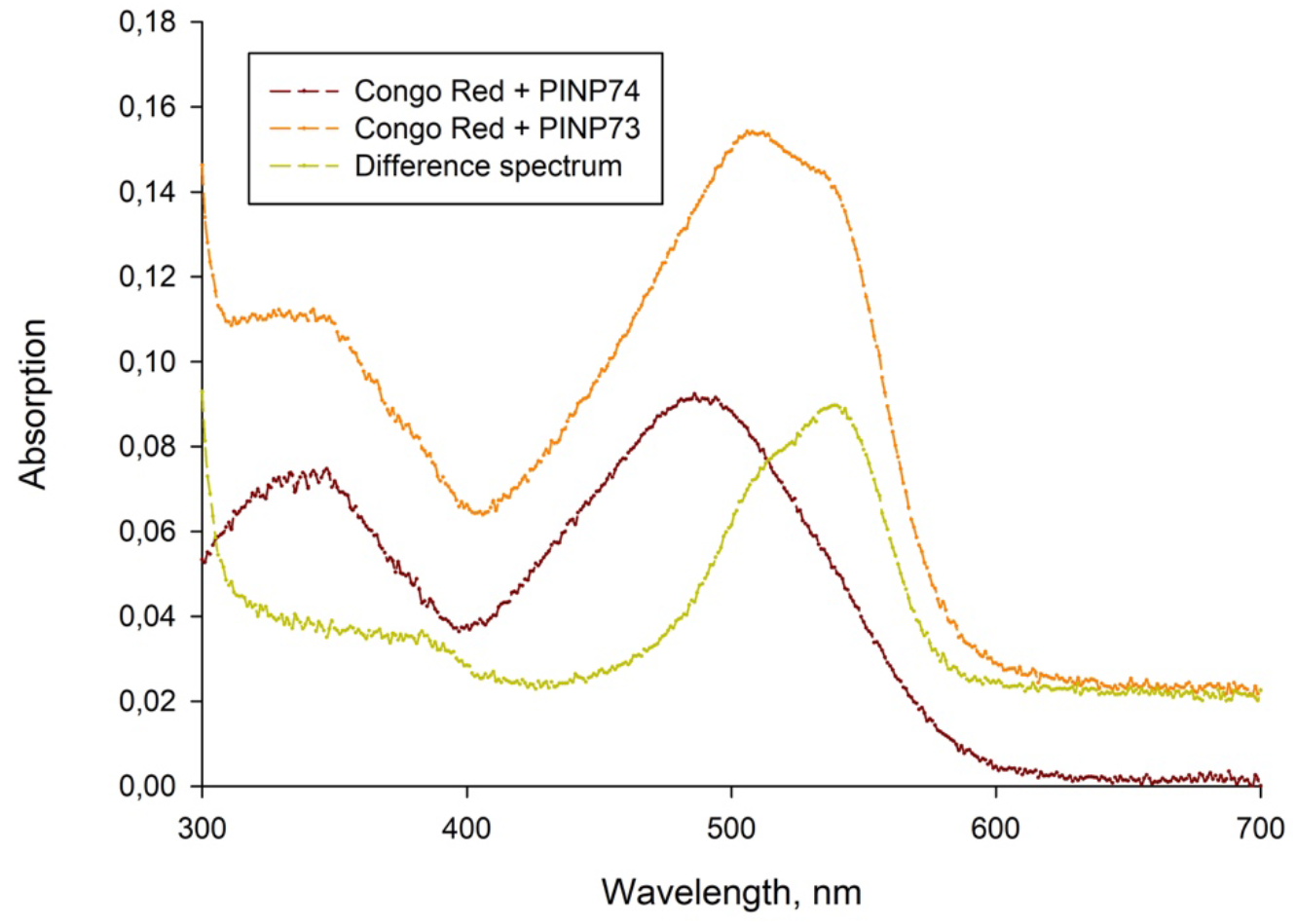
Absorption spectrum of Congo Red with PINP74 (violet), Congo Red with PINP73 (orange), and difference spectrum

## Discussion

There have been previous attempts to predict the amyloidogenic regions of COVID-19 proteins. At [9] the investigators analyzed the known SARS-CoV2 proteins for the presence of amyloidogenic regions. In the cited work, theoretical prediction was performed using the FoldAmyloid, Waltz, and AGGRESCAN software. Interestingly, the PINP74 peptide is located in the region predicted by the FoldAmyloid as amyloidogenic, but does not exhibit amyloidogenicity. Peptide PINP73 is also defined by FoldAmyloid as amyloidogenic, but it also falls within the arches region predicted by the Arches software and is a part of a mirror-symmetric motif. It cannot be said with certainty that the use of these three programs (Arches, FoldAmyloid, and symmetry search) can reliably predict amyloidogenic peptides derived from the whole protein, however, in this work, such an approach made it possible to detect an amyloidogenic peptide. The combination of beta-turn forming proneness, symmetry, and context of amino acid residues may be the key to predicting the ability to form amyloid-like fibrils. In the light of recently appearing in databases of fibril structures obtained using cryo-electron microscopy, such a relationship between the features of the primary structure and the ability to homooligomerization is the subject of our further research. With regard to viruses, in particular SARS-CoV2, the ability of its proteins and protein fragments to amyloidogenesis is unlikely to be an artifact. Interestingly, for the S-protein, the ability of its fragment to form amyloid-like fibrils was confirmed experimentally [27]. Also, the ability of SARS-CoV2 proteins to form amyloid-like fibrils is considered by a number of researchers as one of the possible pathogenicity factors [11], [12].

In this work, we demonstrate for the first time the ability of the NSP7 fragment to form amyloid-like fibrils *in vitro*. Further studies will be aimed at studying the ability of fibrils formed by this peptide to induce coaggregation of the full-length recombinant NSP7 protein and to study the ability of the peptide to act as an antiviral agent in a cell culture model.

## Acknowledgements

The reported study was funded by RFBR, project number 20-04-60491

